# The T4bSS of *Legionella* features a two-step secretion pathway with an inner membrane intermediate for secretion of transmembrane effectors

**DOI:** 10.1101/2024.03.14.584949

**Authors:** Silke Malmsheimer, Iwan Grin, Erwin Bohn, Mirita Franz-Wachtel, Boris Macek, Tobias Sahr, Fabian Smollich, David Chetrit, Amit Meir, Craig Roy, Carmen Buchrieser, Samuel Wagner

**Affiliations:** University of Tübingen, Interfaculty Institute of Microbiology and Infection Medicine (IMIT), Section of Cellular and Molecular Microbiology, Elfriede-Aulhorn-Str. 6, 72076 Tübingen, Germany; Institut de Recherche en Infectiologie de Montpellier, Equipe Kremer, UMR 9004 – CNRS / UM, 1919 route de Mende, 34293 Montpellier cedex 5, France; University of Tübingen, Interfaculty Institute of Microbiology and Infection Medicine (IMIT), Institute of Medical Microbiology and Hygiene, Elfriede-Aulhorn-Str. 6, 72076 Tübingen, Germany; University of Tübingen, Proteome Center Tübingen, Auf der Morgenstelle 15, 72076 Tübingen, Germany; Institute Pasteur, Université Paris Cité, Biologie des Bactéries Intracellulaires, 75015 Paris, France; Yale University, Department of Microbial Pathogenesis, 295 Congress Avenue, New Haven, CT 06536-0812, USA; Birkbeck Institute of Structural and Molecular Biology, Birkbeck and UCL, Malet Street, London WC1E 7HX, UK; University of Glasgow, MRC Centre for Virus Research, School of Infection and Immunity, Glasgow, UK; Excellence Cluster “Controlling Microbes to Fight Infections” (CMFI), Elfriede-Aulhorn-Str. 6, 72076 Tübingen, Germany; German Center for Infection Research (DZIF), partner-site Tübingen, Elfriede-Aulhorn-Str. 6, 72076 Tübingen, Germany

## Abstract

To promote intracellular survival and infection, *Legionella spp.* translocate hundreds of effector proteins into eukaryotic host cells using a type IV b protein secretion system (T4bSS). T4bSS are well known to translocate soluble as well as transmembrane domain-containing effector proteins (TMD-effectors) but the mechanisms of secretion are still poorly understood. Herein we investigated the secretion of hydrophobic TMD-effectors, of which about 80 were previously reported to be encoded by *L. pneumophila*. A proteomic analysis of fractionated membranes revealed that TMD-effectors are targeted to and inserted into the bacterial inner membranes of *L. pneumophila* independent of the presence of a functional T4bSS. While the T4bSS chaperones IcmS and IcmW were critical for secretion of all tested TMD-effectors, they did not influence inner membrane targeting of these proteins. As for soluble effector proteins, translocation of TMD-effectors into host cells depended on a C-terminal secretion signal and this signal needed to be presented towards the cytoplasmic side of the inner membrane. A different secretion behavior of TMD- and soluble effectors and the need for small periplasmic loops within TMD-effectors provided strong evidence that TMD-effectors are secreted in a two-step secretion process: Initially, an inner membrane intermediate is formed, that is extracted towards the cytoplasmic side, possibly by the help of the type IV coupling protein complex and subsequently secreted into eukaryotic host cells by the T4bSS core complex. Overall, our study highlights the amazing versatility of T4bSS to secrete soluble and TMD-effectors from different subcellular locations of the bacterial cell.

## Introduction

In Gram-negative bacteria several different secretion machines have been described that export protein substrates from the bacterial cytoplasm in one step across the entire cell envelope, comprising the cytoplasmic and inner membrane (IM), the periplasm and the outer membrane (OM) [1]. Type III, type IV, and type VI secretion systems (T3SS, T4SS, T6SS) possess the additional feature of translocating substrate proteins, so called effectors, directly into eukaryotic target cells [2]. *Legionella spp.* employ a T4bSS, also called Dot/Icm (defective for organelle trafficking/intracellular multiplication defect) secretion system to establish a parasitic lifestyle within vacuoles of amoeba. To this end, *Legionella* translocate a cocktail of more than 300 different effector proteins via the T4bSS that subtly manipulate host cell biology for the benefit of *Legionella* [3–5]. The same mechanisms are used by human pathogenic *Legionella* spp. like *L. pneumophila* to infect alveolar macrophages, leading to Legionnaires’ disease. Evolutionary and structurally related to bacterial conjugation machines, the T4bSS of *Legionella* comprises at least 32 different structural components of which most are critical for secretion function of this molecular machine [6–9]. The T4bSS consists of a core complex spanning the entire bacterial cell envelope and of a type IV coupling complex (T4CC) in the inner membrane. While the core complex is thought to form the actual ATP-fueled secretion machine, the T4CC seems to serve as the primary receptor for substrate proteins [10].

The targeting signal for T4bSS resides at the C-terminus of substrate proteins and is about 20-30 residues long [11,12]. Although the C-termini of T4bSS substrates do not share detectable sequence homology, they do appear to be enriched in certain residues including small, polar and/or charged amino acids, suggesting a related motif or structure [13]. For a number of effectors, a so-called E-block motif has been identified at the C-terminus that consists of several consecutive glutamic acid residues [14]. Moreover, the targeting of some T4bSS substrates is mediated by the T4bSS chaperones IcmSW and LvgA that seem to act as adapters between effectors and the T4CC [7,15]. It was proposed that E-block motif-containing effectors do not require IcmSW for targeting to the secretion machine but directly bind to DotM, an IM protein of the TC4P with a sizable, positively charged cytoplasmic domain [16]. From the T4CC effectors are believed to be handed over to the ATPases of the system that subject them into the secretion channel.

About 25% of all effectors of T4bSS are transmembrane domain-containing effectors (TMD-substrates) that have their final destination in one of the membranes of the eukaryotic host [17–19]. Transmembrane segments (TMSs) within effectors pose a potential targeting conflict as two sequential secretion signals for two potentially incompatible pathways are concatenated in the same protein: the C-terminal secretion signal for transport through the T4bSS on the one hand and hydrophobic TMS that signal inner membrane targeting on the other hand [20]. We recently investigated how targeting discrimination between these membrane protein targeting pathways is achieved for TMD-effectors of the T3SS inside *S.* Typhimurium [19]. A balanced hydrophobicity of the TMS of T3SS substrates is one factor supporting targeting discrimination: While being sufficiently hydrophobic for principal membrane integration, inner membrane targeting was generally not facilitated by these segments. Also, we reported that the binding of the hydrophobic TMS of the effector SseF by its cognate T3SS chaperone prevented erroneous inner membrane targeting and integration.

Also a subset of TMD-substrates of T4bSS of *L. pneumophila* possess TMS of reduced hydrophobicity and by this avoid SRP-targeting and facilitate direct delivery of these effectors to the T4bSS machinery from the cytoplasmic side [19]. In addition, T4bSS are able to secrete TMD-effectors of higher hydrophobicity [17,19] that could be targeted by the signal recognition particle (SRP) and integrate into the bacterial inner membrane in a co-translational fashion. This would likely occur unhindered from T4bSS targeting mechanisms as the main targeting signal of T4bSS effectors reside at their C-terminus. Consequently, these TMD-effectors may need to be accepted by the T4bSS from within the bacterial inner membrane and thus be secreted by a unique two-step secretion pathway.

In this study, we elucidated whether TMD-effectors with higher hydrophobicity are targeted to and inserted into the bacterial inner membrane prior to recruitment to the T4bSS machinery or whether they can avoid inner membrane targeting by chaperone binding similar to TMD-effectors of the T3SS. Using membrane fractionation as well as urea extraction, we could show that TMD-effectors of *L. pneumophila* can indeed be found properly integrated in the bacterial inner membrane. The same results were obtained when the type IV chaperones IcmSW were overexpressed, suggesting that these chaperones do not contribute to avoiding the inner membrane targeting of TMD-effectors. Investigation of the membrane topology of TMD-effectors showed that they largely possess a N_in_-C_in_ topology in the bacterial inner membrane with only small loops of few amino acids located in the periplasm. Similar to soluble effectors, the C-terminal translocation signal of TMD-effectors needed to be located in the cytoplasm for successful secretion and possible internal signals as well as the presence of the chaperones IcmSW were crucial for the successful translocation of TMD-effectors into host cells. Based on these results we propose that TMD-effectors in *L. pneumophila* follow a two-step secretion pathway with an inner membrane intermediate, followed by recognition by components of the T4CC and subsequent secretion through the T4bSS core complex from the cytoplasmic side.

## Results

### T4bSS TMD-effectors are integrated into the inner membrane of *L. pneumophila*

T4bSS are able to secrete TMD-effectors of higher hydrophobicity [17,19]. These proteins are expected to be recognized by the SRP and integrate into the bacterial inner membrane in a co-translational fashion, unaffected by potential T4bSS targeting, as the main targeting signal of T4bSS effectors resides at the C-terminus.

To evaluate the integration of TMD-effectors into the inner membrane of *L. pneumophila* experimentally, we analyzed the localization of T4bSS-effectors across a membrane-fractionating sucrose gradient after sucrose density equilibrium centrifugation. Crude membranes were separated in a continuous 30-53% sucrose gradient and subsequently fractionated into 12 fractions of equal volume. The fractions were analyzed by mass spectrometry. Similar to membrane fractionation of *Escherichia coli* and *Salmonella* Typhimurium [21,22], cytoplasmic proteins peaked in the low density fractions one and two, while outer membrane proteins were mainly found in the high density fractions nine and ten (Fig. 1A). Inner membrane proteins showed a more dispersed fractionation profile with a slight peak in fractions three and four. 26 T4bSS TMD-effectors and 54 soluble effectors could be detected and quantified by mass spectrometry. TMD-effectors exhibited a fractionation profile similar to reported inner membrane proteins (Fig. 1A). In contrast, one subset of soluble effectors fractionated like cytoplasmic proteins while another subset fractionated like inner membrane proteins. The mass spectrometry results were complemented by Western blotting and immunodetection of two plasmid-expressed, HA-epitope tagged TMD-effectors (^HA^LegC2 and ^HA^LegC3) and one soluble effector (^HA^SidM) across the twelve fractions. Also in this analysis, the fractionation profiles of ^HA^LegC2 and ^HA^LegC3 resembled the ones of reported inner membrane proteins while the fractionation profile of ^HA^SidM compared to the ones of cytoplasmic proteins (Fig. 1B). Quantification of the same un-tagged endogenously expressed proteins by mass spectrometry showed essentially the same distribution in the sucrose gradient (Fig. 1C). In the absence of the T4bSS, no substantial change in running behavior of these effectors was observed (Fig. 1C), indicating that inner membrane targeting is the dominant factor contributing to the subcellular localization of effector proteins.

**Fig. 1.**
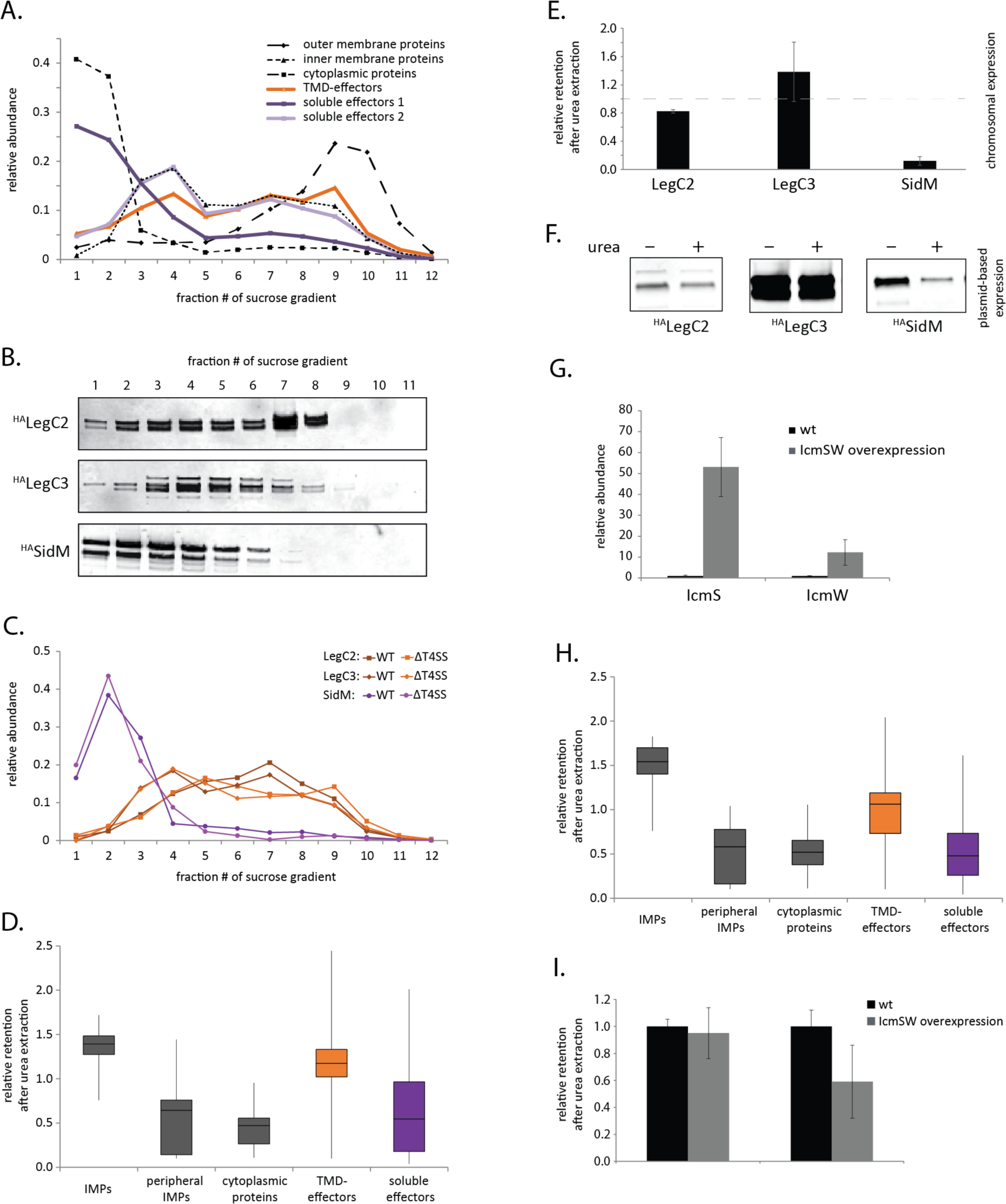
Analysis of the subcellular localization of TMD-effectors in *L. pneumophila.* **A.** The relative abundance of the indicated effector classes in 12 fractions of a membrane-fractionating sucrose gradient as analyzed by mass spectrometry. For reference, reported cytoplasmic, inner membrane and outer membrane proteins were plotted. Substrates of the T4bSS system were found by matching the proteins identified by mass spectrometry to the Secret4 database [23]. The data points represent the mean of at least three independent experiments. **B.** The accumulation levels of the indicated plasmid-expressed and epitope-tagged effector proteins in 12 fractions of a membrane-fractionating sucrose gradient were analyzed by SDS-PAGE, Western blotting and immunodetection. **C.** The relative abundance of the indicated endogenously expressed effectors in 12 fractions of a membrane-fractionating sucrose gradient as analyzed by mass spectrometry. **D.** The relative retention of the indicated protein groups in urea-extracted inner membrane fractions (2-6 in A) of *L. pneumophila* wild type bacteria as analyzed by mass spectrometry. The mean of three experiments is presented in a box plot. **E.** The relative retention of the indicated endogenously-expressed effectors in urea-extracted inner membrane fractions (2-6 in A) of *L. pneumophila* wild type bacteria as analyzed by mass spectrometry. **F.** The relative retention of the indicated plasmid-expressed and epitope-tagged effector proteins in urea-extracted inner membrane fractions (2-6 in A) of *L. pneumophila* wild type bacteria as analyzed by mass spectrometry. **G.** The relative abundance of IcmS and IcmW in inner membrane fractions of wild type and IcmS and IcmW-overexpressing *L. pneumophila*, respectively, as analyzed by mass spectrometry. **H.** The relative retention of the indicated protein groups in urea-extracted inner membrane fractions (2-6 in A) of *L. pneumophila* overexpressing the T4bSS chaperones IcmS and IcmW as analyzed by mass spectrometry. The mean of three experiments is presented in a box plot. **I.** The relative retention of the indicated endogenously-expressed effectors in urea-extracted inner membrane fractions of wild type and IcmS and IcmW-overexpressing *L. pneumophila* as analyzed by mass spectrometry. p-values are provided in S5 Table.

While cytoplasmic proteins, peripheral inner membrane proteins and soluble effectors could be extracted from the inner membranes by treatment with 8 M urea, TMD-effectors remained associated with inner membranes, similar to integral inner membrane proteins (Fig. 1DE). These results were confirmed for ^HA^LegC2 and ^HA^LegC3 as well as for the soluble effector ^HA^SidM by Western blotting and immunodetection (Fig. 1F). Together these data suggest that TMD-effectors are truly embedded in the inner membrane.

Since we learned that cognate chaperones could prevent erroneous targeting of TMD-effectors of T3SS to the bacterial inner membrane [19], we asked whether the reported T4bSS chaperones IcmS and IcmW also effect the inner membrane localization of TMD-effectors. Thus, we overexpressed IcmS and IcmW simultaneously from an IPTG-inducible plasmid and analyzed the retention of proteins in inner membrane fractions after treatment with 8 M urea like before. 50-fold and 20-fold higher accumulation levels of IcmS and IcmW, respectively (Fig. 1G), did not significantly change the localization of TMD-effectors to the inner membranes (Fig. 1HI), suggesting that these chaperones do not shield TMSs from membrane targeting as it is the case in T3SS. Interestingly, in the sucrose gradient IcmS and IcmW cofractionated with the T4CC and did not behave like soluble cytoplasmic proteins (S1 Fig.). This supports the notion that these two chaperones act like adaptor proteins bound to the T4CC [7] rather than like free cytoplasmic chaperones of the T4bSS.

In summary, membrane fractionation of *L. pneumophila* showed that TMD-effectors are properly integrated into the bacterial inner membrane. This suggests that TMD-effectors follow a two-step secretion pathway with an inner membrane intermediate unless they exhibit a strong potential of mistargeting to the bacterial inner membrane.

### A C_in_ topology facilitates TMD-effector translocation

To be secreted by the T4bSS, inner membrane-integrated TMD-effectors need to be recognized by the secretion machinery. Since recognition of soluble effectors was reported to be accomplished by the T4CC at the cytoplasmic side of the membrane, we reasoned that TMD-effectors might be recognized similarly with a C-terminal T4bSS secretion signal facing the cytoplasm (C_in_). We predicted the topology of the 82 previously reported TMD-effectors using DeepTMHMM [24]. Surprisingly, DeepTMHMM picked up only 27 of them as transmembrane proteins (S1 Table, S2 Table, S3 Table), which may result from the fact that DeepTMHMM was not trained on TMD-effectors injected by bacterial pathogens. Of the 27 TMD-effectors that were picked up by DeepTMHMM, 21 (78%) were predicted to exhibit an N_in_/C_in_ topology (Fig. 2A) and 22 (81%) were predicted to possess a transmembrane topology of low complexity with only one or two TMS (Fig. 2B).

**Fig. 2.**
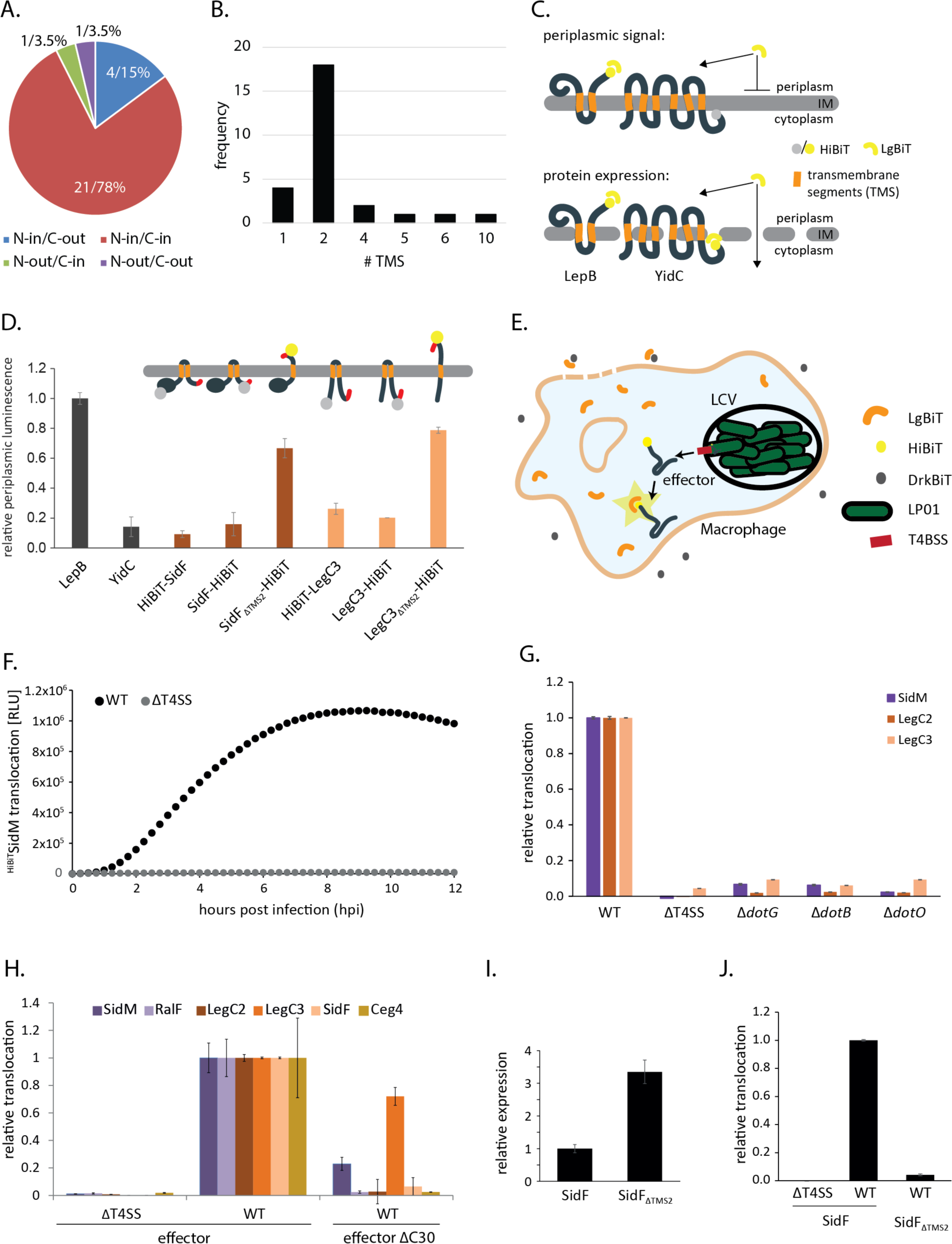
Analysis of the relevance of the presence and localization of the C-terminal signal for TMD-effector translocation. **A.** Pie-chart showing the proportion of the indicated transmembrane topologies of the TMD-effectors picked up by DeepTMHMM. **B.** Frequency of the number of transmembrane segments in the TMD-effectors picked up by DeepTMHMM. **C.** Illustration of the principle of the split NanoLuc-based topology assay. HiBiT fused to a terminus of the protein of interest (here topology control proteins LepB and YidC) is complemented by the addition of LgBiT and luminescence is assayed. The transmembrane topology is deduced from the periplasmic luminescence (top) relative to the total luminescence in lysed bacteria (bottom) **D.** Relative periplasmic luminescence of the indicated HibiT-fusion proteins, assayed as shown in C. The data points represent the mean (± standard deviation) of at least three independent experiments. **E.** Illustration of the principle of the split NanoLuc-based translocation assay. HibiT fused to the N-terminus of a translocated effector is complemented by LgBiT stably expressed in the cytoplasm of RAW 264.7 macrophages and luminescence is assayed. Extracellular LgBiT resulting from cell lysis is quenched by the addition of DrkBiT. F. Relative translocation of plasmid-expressed HiBiT-SidM into RAW 264.7 macrophages as assayed over time in a plate reader using the split-NanoLuc translocation assay. **G.** Relative translocation of the indicated effectors into RAW 264.7 macrophages using *L. pneumophila* wild type and the indicated mutants. The measurement was performed 4 h post infection by the split NanoLuc-based translocation assay shown in E. **H.** Relative translocation of the indicated effectors w/ and w/o C-terminal secretion signal in the wild type and T4bSS-deficient mutant bacteria. The measurement was performed 4 h post infection by the split NanoLuc-based translocation assay shown in E. **I.** Relative expression of SidF w/ and w/o its second TMS in the wild type and T4bSS-deficient mutant bacteria. **J.** Relative translocation of SidF w/ and w/o its second TMS in the wild type and T4bSS-deficient mutant bacteria. The data points represent the mean (± standard deviation) of at least three independent experiments. p-values are provided in S5 Table.

To validate the C_in_-transmembrane topology of selected TMD-effectors within *L. pneumophila*, we developed a split-NanoLuc based topology assay [25]. Split NanoLuc utilizes bi-molecular complementation of NanoLuc luciferase, comprising a 18 kDa fragment of NanoLuc, LgBiT, and a 1.3 kDa peptide, HiBiT, which has a very high affinity for LgBiT and complements its luciferase function [25]. The transmembrane topology was assessed by testing the accessibility of HiBiT fused to the N-or C-termini of the proteins of interest. HiBiT localized at the periplasmic side of the membrane was expected to be accessible to LgBiT added to EDTA-treated *L. pneumophila*, thus giving rise to a luminescence signal. In contrast, HiBiT localized at the cytoplasmic side of the membrane should be inaccessible for LgBiT unless the inner membrane is permeabilized (Fig. 2C). The assay was validated by assessment of the transmembrane topology of *L. pneumophila* homologs of the well-studied integral membrane proteins LepB and YidC [26,27] (Fig. 2D). Both proteins showed the expected topology: C_out_ for LepB and C_in_ for YidC. The TMD-effectors SidF and LegC3, respectively, proved to exhibit a N_in_-C_in_ topology unless their second TMS was deleted, which resulted in a change to a N_in_-C_out_ topology (Fig. 2D).

To further elucidate the importance of the presence and localization of the C-terminal secretion signal of TMD-effectors for their translocation into host cells, we adapted a split-NanoLuc based translocation assay [28–30]. Plasmid-expressed HiBiT-effector-fusions were translocated by *L. pneumophila* into RAW 264.7 macrophages, stably expressing LgBiT in their cytoplasm and luminescence was assayed over time in a microtiter plate reader. The peptide DrkBiT, which exhibits a very high affinity for LgBiT but does not complement luminescence activity [31], was introduced into the infection buffer to effectively suppress any luminescence signals that could potentially emerge from extracellular LgBiT after macrophage cell lysis (Fig. 2E). Using this set-up, we could follow the kinetics of effector translocation over several hours post infection as exemplified for a HiBiT-SidM fusion in Fig. 2F. Translocation of effectors proved to be strongly dependent on a functional T4bSS as deletion of the entire system or deletion of the structural component DotG or the ATPases DotB or DotO, respectively, led to a reduction of the luminescence signal of >90% for the tested effectors SidM, LegC2 and LegC3 (Fig. 2G). Like most previously reported soluble T4bSS effectors, three of the four selected TMD-effectors as well as the soluble effectors SidM and RalF could not be translocated without their C-terminal 30 amino acids (Δc30) (Fig. 2H). Only the translocation of the TMD-effector LegC3 reached about 70% of the wild type levels 4 hours post infection in the absence of its C-terminus. We then tested whether the cytoplasmic localization of the C-terminal signal was important for the efficient translocation of the TMD-effector SidF. Deletion of its second TMS, resulting in a periplasmic presentation of the C-terminus as shown in Fig. 2D, almost completely abrogated the translocation of SidF into host cells, despite the more than 3-fold higher expression levels of this mutant (Fig. 2IJ). Together these results provide strong evidence that TMD-effectors are also recognized by C-terminal T4bSS signals and that these signals must be presented to the cytoplasmic side of the bacterial membrane to support successful secretion. Thus, recognition of the C-terminal signals of TMD-effectors may also occur by the T4CC in a manner analogous to the recognition of soluble effector proteins.

### TMD-effectors are recognized by IcmSW

As introduced above, it was proposed previously that T4bSS effector proteins harboring an E-block motif at their C-terminus are recognized by the T4CC component DotM but do not require the aid of the chaperones IcmS and IcmW. Effectors lacking an E-block motif instead, often require these two chaperones [16]. Of the 82 previously reported TMD-effectors of *L. pneumophila* LP01 [19], 65% exhibit a clear E-block motif while 35% do not (Fig. 3A). We used the split-NanoLuc translocation assay to assess the relevance of IcmS and IcmW as well as of DotM for translocation of TMD-effectors into RAW 264.7 macrophages depending on their E-block motif.

**Fig. 3.**
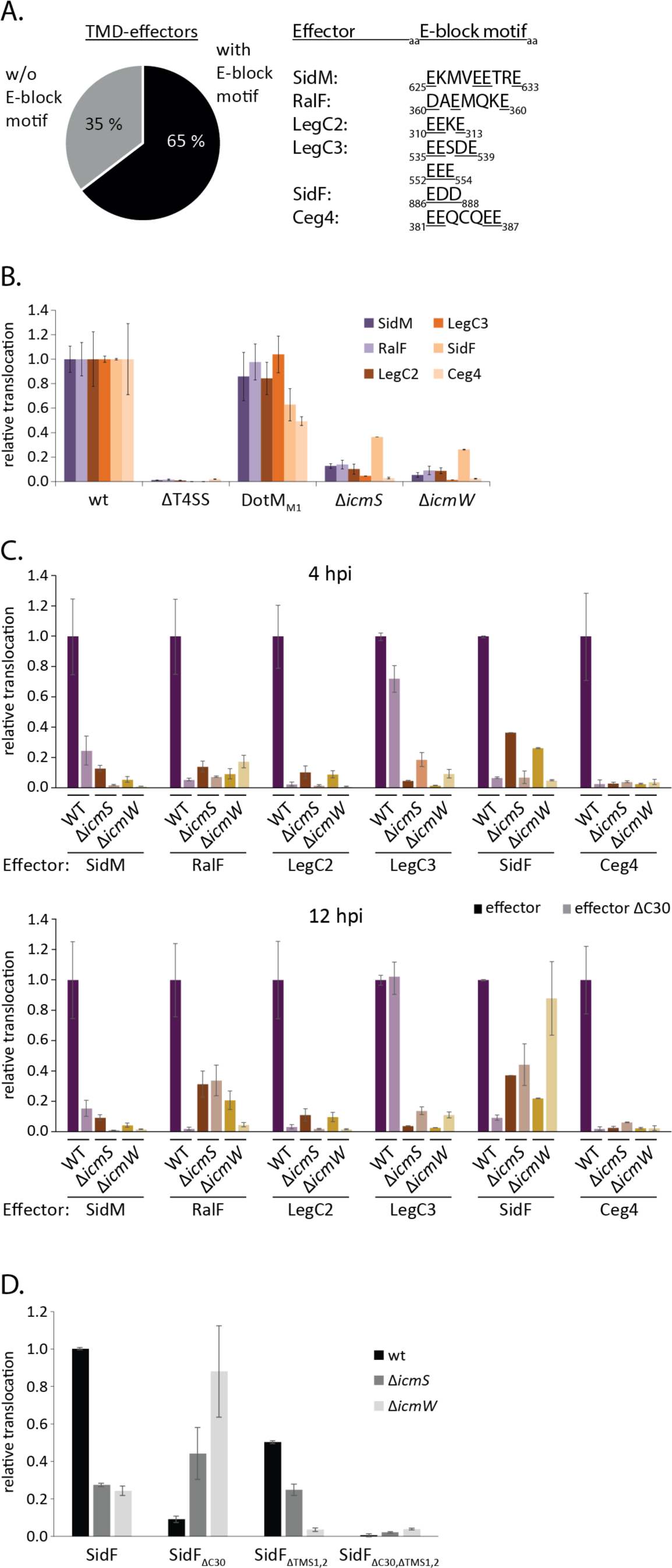
Analysis of TMD-effector translocation in mutants of the T4CC. **A.** Analysis of the C-terminal 30 amino acids of all previously reported TMD-effectors for the presence of an E-block motif according to the classification of Huang *et al.* [14]. **B.** Relative translocation of indicated effectors in the wild type bacteria as well as in a DotM mutant (DotM1) with decreased ability to bind Glutamine-rich peptides [16] and in strains lacking *icmS*, *icmW*, respectively. The measurement was performed 4 h post infection by the split NanoLuc-based translocation assay shown in Fig. 2E. The data points represent the mean (± standard deviation) of at least three independent experiments. **C.** As in B but showing indicated effectors w/ and w/o C-terminal secretion signal in wild type, T4bSS- and *icmS* or *icmW*-deficient mutant bacteria. The Luminescence signal after 4 hours post infection (upper panel) or 12 hours post infection (lower panel) is displayed. **D.** As in C but showing SidF w/ and w/o its second TMS and w/ and w/o its C-terminal secretion signal. The luminescence signal 12 hours post infection is displayed. p-values are provided in S5 Table.

Surprisingly, the DotM_M1_ mutant (R196E, R197E), reported to block translocation of substrates containing E-block motives [16], did not lead to a decrease in the translocation efficiency of SidM, RalF, LegC2 and LegC3 despite the presence of an E-block motif within the C-termini of these effectors (Fig. 3B, S3 Table). This result is most noticeable for LegC3 which even has two strong E-block motifs, EESDE and EEE (Fig. 3A). The translocation of SidF and Ceg4 was reduced by 40% and 50% of the wild type levels, respectively, in the DotM_M1_ mutant.

In Δ*icmS* and Δ*icmW* single mutants, both soluble effectors SidM and RalF as well as the TMD-effectors LegC2, LegC3 and Ceg4 showed a >80% reduction in translocation (Fig. 3B). Only SidF showed a residual translocation of up to 40% in the chaperone deletion mutants (Fig. 3B). Simultaneous deletion of one of the T4bSS chaperones and the C-terminal secretion signal resulted in different translocation behaviors (Fig. 3C). After four hours of infection, a synergistic reduction of effector translocation in the double mutants was observed for SidM, LegC2 and SidF. In the case of Ceg4, translocation levels were very low in all tested mutants. In contrast, for LegC3, simultaneous deletion of the C-terminal 30 amino acids and *icmS* or *icmW*, respectively, resulted in a higher translocation than deleting *icmS* or *icmW* alone (Fig. 3C). Remarkably, translocation of SidF was restored to 40% and 60%, respectively, in the *icmS*/ΔC30 and *icmW*/ΔC30 double mutants after 12 hours of infection (Fig. 3C). The restoring effect of these double mutants was only observed for membrane localized SidF. When we deleted both TMS in addition, resulting in a soluble SidF, no translocation could be observed in the absence of the C-terminal 30 amino acids at all despite a pronounced translocation of the ΔTMS mutant in the presence of the C-terminus (Fig. 3D).

Overall, these data show that the chaperones IcmS and IcmW play a strong role in the translocation process of TMD-effectors, similar to what has been reported for soluble effector proteins. However, the example of SidF demonstrates that the simultaneous deletion of IcmS or IcmW and the C-terminal secretion signal can have a different effect on translocation depending on the presence or absence of TMS in the effector and that a delayed translocation is possible in the absence of both, the chaperones and the C-terminal signal as long as the effector contains TMS. These differential observations for soluble vs. TMD-SidF strongly support the notion that SidF follows a two-step secretion process with an inner membrane intermediate.

### TMD-effectors exhibit short periplasmic loops

Once integrated into the bacterial inner membrane and recognized by the Dot/Icm machinery, TMD-effectors need to be extracted from the lipid bilayer to be handed over to the core complex of the T4bSS for subsequent translocation. ATPases that could mediate extraction from the membrane reside on its cytoplasmic side, thus extraction of TMD-effectors towards the cytoplasm is most likely.

The DeepTMHMM topology prediction revealed that the majority of TMD-effectors picked up by this program have only very short periplasmic loops with a length of less than 30 amino acids (Fig. 4A). To assess if bigger protein domains located in the periplasm might interfere with protein translocation by preventing extraction from the membrane towards the cytoplasm, more extensive protein domains were inserted into the periplasmic loop of SidF (Fig. 4B). Luminescence measurements showed that the insertion of the 11 residues long HiBiT peptide resulted in a slight decrease in SidF translocation whereas the fusion of SidF with the weakly-folding 8.6 kDa ubiquitin_E3G,E13G_ variant resulted in a 80% decrease in translocation (Fig. 4C). The ubiquitin_E3G,E13G_ variant was shown to be translocation-permissive in T4bSS and T3SS, previously [32,33], suggesting that the failure of translocation of the SidF-ubiquitin_E3G,E13G_ variant results from the inability to extract this SidF variant from the membrane. Protein expression and inner membrane localization remained unaffected in these SidF variants (Fig 4D).

**Fig. 4.**
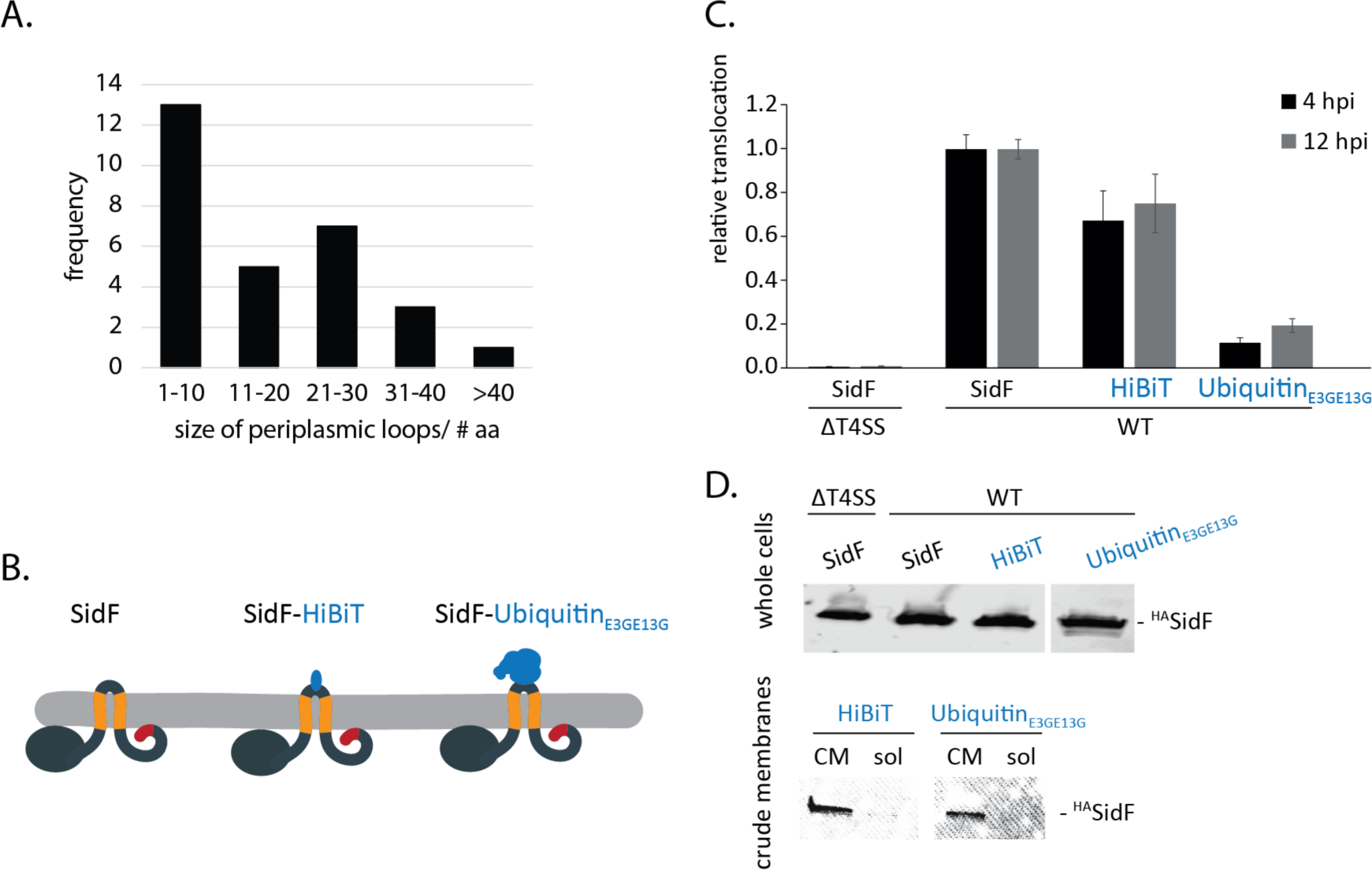
Role of the size of the periplasmic loop of SidF in translocation. **A.** Frequency of the size of periplasmic loops in the TMD-effectors picked up by DeepTMHMM. **B.** Illustration of SidF-fusion proteins embedded in the bacterial inner membrane. **C.** Relative translocation of the indicated SidF-fusion proteins in the wild type bacteria. The measurement was performed 4 and 12 h post infection by the split NanoLuc-based translocation assay shown in Fig. 2E. The data points represent the mean (± standard deviation) of at least three independent experiments. **D.** Western blot analysis of whole cells, crude membrane (CM) and soluble fractions of *L. pneumophila* expressing the indicated HA-epitope-tagged SidF-fusion proteins. p-values are provided in S5 Table.

These data provide evidence that larger periplasmic domains of TMD-effectors compromise their translocation, possibly by interfering with the ability to extract these effectors from the inner membrane towards the cytoplasmic side.

## Discussion

20-30% of all proteins secreted by the T4bSS of *Legionella pneumophila* are transmembrane proteins that presumably fulfill their function within membranes of the eukaryotic host. Although the functional role of some of these TMD-effectors has been elucidated in previous studies, the question of how they can be targeted to and recognized by the secretion system within *Legionella* has not been clarified to date. Here we provide evidence that many TMD-effectors of *L. pneumophila* follow a unique two-step secretion mechanism with an inner membrane intermediate before T4bSS-dependent translocation into the host cell.

Unlike soluble effectors, TMD-effectors contain hydrophobic segments that are needed for successful integration into the lipid bilayer of their target membrane. Inside the bacterium, these TMS are in general co-translationally recognized by the SRP, which consequently leads to the integration of the protein into the bacterial inner membrane. In a previous study, we were able to show that intrinsic properties of TMD-effectors of T3SSs such as a balanced hydrophobicity of the TMS or binding by cognate T3SS chaperones results in avoidance of inner membrane targeting and insertion of these effector proteins [19]. This is critical for T3SS, which can only accept substrates that are directly targeted to the system in the cytoplasm. In contrast, T4SSs have also been shown to secrete substrate proteins from the periplasm as exemplified by the *B. pertussis* toxin as well as effector proteins of the VirB T4SS of *Brucella* spp. These effectors were shown to be translocated across the bacterial inner membrane using the Sec system in a first step, and then they cross the bacterial outer membrane in a T4SS-dependent manner in a second step [34–36]. Also, many TMD-effectors of *L. pneumophila* exhibit a rather high hydrophobicity within their TMS [19], making their recognition by the SRP and consequently an inner membrane secretion intermediate very likely. Using membrane fractionation by a sucrose gradient centrifugation protocol, as well as urea extraction, we show that TMD-effectors in *L. pneumophila* can indeed be found properly integrated into the bacterial inner membrane.

To prevent inner membrane targeting, the *Salmonella* Typhimurim T3SS substrate SseF that contains TMS of stronger hydrophobicity was shown to require the action of its cognate chaperone SscB. It was hypothesized that a co-translational binding of the chaperone to the effector, also to the hydrophobic TMS, prevents the co-translational recognition and erroneous targeting of SseF by the SRP [19]. Thus, T3SS may functionally combine co-translational targeting of substrates and post-translational secretion. In contrast to substrates of T3SS, substrates of T4SS harbor C-terminal secretion signals and very ill-defined binding sites for the chaperones of the system. Thus, a co-translational prevention of inner membrane targeting by T4SS chaperones is highly unlikely. Our herein presented results support this notion. Although, translocation of all investigated TMD-effectors was shown to be IcmSW-dependent, overexpression of these chaperones did not affect the inner membrane integration of TMD-effectors (Fig. 1HI). Our data even suggest that the functionally relevant proportion of IcmSW is membrane associated and does not exist in a free, cytoplasmic form, which would be the form to interfere with membrane targeting of TMD-effectors. Thus, our data support the notion that IcmSW mainly act as adaptor proteins between T4SS substrates and the T4CC.

Together these results provide strong evidence that T4bSS-secreted membrane proteins are exported by a two-step mechanism with an inner membrane intermediate. In contrast to other protein secretion systems of bacteria, the T4bSS system of *L. pneumophila* secretes over 300 different proteins of which 20-30% are predicted as membrane proteins. It is conceivable that the inner membrane is used as an intracellular storage of T4bSS-secreted TMD-effectors that serves to prevent their aggregation until secretion into the host cell.

Once inserted into the bacterial inner membrane, TMD-effectors need to be recognized by the T4bSS machinery for subsequent translocation into the host cell. Our results suggest that TMD-effectors are recognized by the T4CC at the cytoplasmic side of the bacterial inner membrane as they do tend to exhibit a C_in_-topology. However, the T4CC is a large platform with different effector docking sites and the TMD-effectors may be recognized differently than their soluble counterparts. Moreover, the cytoplasmic presentation of the C-terminal signal seems to be crucial for successful effector translocation. After inner membrane insertion, TMD-effectors need to access the core complex’ secretion channel. This could be achieved by lateral access in the plane of the membrane or after membrane extraction of the TMD-effectors towards the periplasmic or cytoplasmic side. As the T4bSS core secretion channel is fenced by TMDs, a lateral access is unlikely [9]. Extraction of transmembrane proteins from the lipid bilayer is highly energy-consuming. The small membrane-integral part of TMD-effectors as well as the small periplasmic loops may be necessary for their extraction towards the cytoplasmic face of the membrane prior to effector translocation. Based on the notion that TMD-effectors most probably are recognized by DotL-bound IcmSW, we hypothesize that DotL might provide the energy for effector extraction from the bacterial inner membrane by ATP hydrolysis.

The involvement of the T4CC in effector recognition was already shown previously. It was reported that effectors are either bound by IcmSW [37] or that they are IcmSW-independent by harboring a negatively charged E-block motif within their C-terminal T4bSS signal that interacts with the positively charged surface of DotM [16]. It was also stated that effectors with a strong secretion signal consisting of a large glutamic acid stretch and several hydrophobic residues at the C-terminal end of the effector are less IcmSW-dependent than effectors with a weak signal. In contrast, we show herein that all six investigated soluble as well as TMD-effector proteins did not only show an IcmSW-dependent secretion phenotype but also harbor an E-block motif within their C-terminal signal. Interestingly, disrupting identified interaction sites in DotM [16] by a point mutation did not interfere with effector translocation, indicating that the presence of an E-block motif does not necessarily result in the anticipated recognition of the effector protein by DotM. Which other components of the T4CC are involved in the recognition of the C-terminal secretion signal or if the secretion signal is only important at a later point in the secretion mechanism has yet to be elucidated.

Surprisingly, simultaneous deletion of *icmS* or *icmW* and the C-terminal secretion signal led to an increased secretion of LegC3 after 4 h of infection and of SidF after 12 h of infection compared to the respective single deletion mutants. In the case of SidF, this phenomenon was specific for the membrane-localized protein and absent in case of a soluble variant lacking the TMS. On the one hand, these differential secretion requirements for the membrane-localized and soluble SidF, respectively, support the notion that SidF is indeed secreted with an inner membrane intermediate. On the other hand, these results suggest that IcmS and IcmW and the C-terminal signal of TMD-effectors function synergistically and are critical for the efficient translocation of TMD-effectors but that they are not essential for principal secretion. The C-terminus of DotL that binds IcmS and IcmW is highly dynamic and flexible. It may allow T4CC-bound IcmS and IcmW to reach to the inner membrane and to utilize ATP hydrolysis by DotL for retraction and membrane extraction of TMD-effectors.

In summary, we provide substantial evidence that TMD-effectors of the T4bSS of *L. pneumophila* follow a two-step secretion pathway with an inner membrane intermediate, revealing a novel aspect of the versatility of secretion capabilities of these molecular secretion machines (Fig. 5). T4SS are the only known bacterial protein secretion systems that are secreting soluble substrates from the cytoplasm and the periplasm as well as integral inner membrane proteins. It remains to be elucidated in the future, what the exact functional mechanisms for any of these different secretion modes are.

**Fig. 5.**
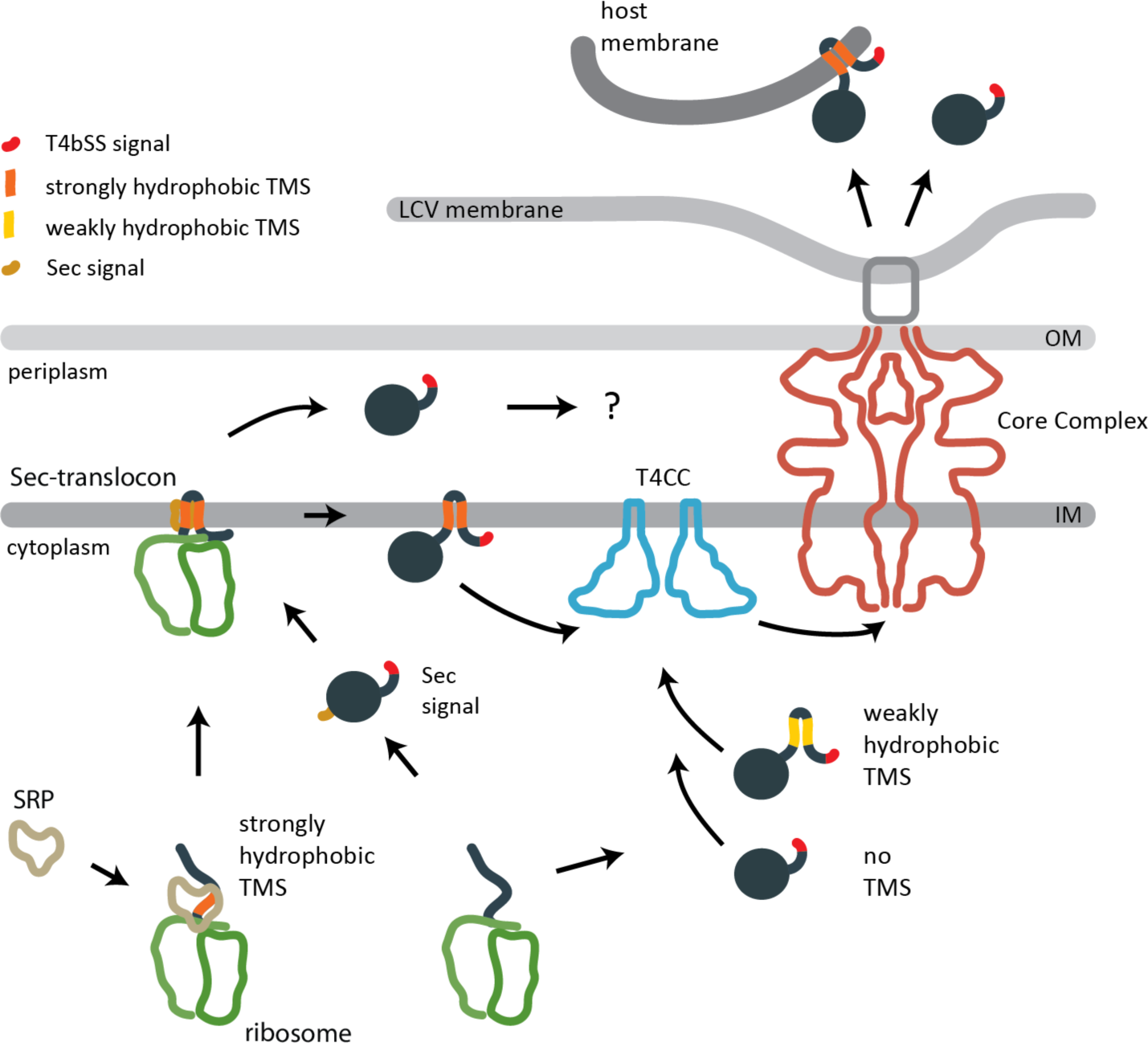
Model of the secretion pathways of T4bSS effectors in *L. pneumophila.* T4bSS are able to secrete soluble and TMD-effectors into eukaryotic host cells. Soluble effectors and weakly hydrophobic TMD-effectors may be targeted directly to the type 4 coupling complex (T4CC). More strongly hydrophobic TMD-effectors may be targeted by the signal recognition particle (SRP) to the Sec translocon and subsequently inserted into the bacterial inner membrane. We propose that they are then recognized by the T4CC based on their C-terminal T4bSS signal, extracted towards the cytoplasmic side of the membrane and subsequently secreted by the T4bSS core complex. Small periplasmic loops may facilitate their extraction from the membrane. A low complexity of the TMD may facilitate the extraction from the bacterial inner membrane as well as the insertion into the host membrane. Soluble effectors harboring a Sec secretion signal may be targeted to the Sec translocon and released into the periplasm. Here they may be recognized by the T4bSS core complex directly and secreted subsequently. Four of the previously listed TMD-effectors were now predicted to contain cleavable Sec signals by DeepTMHMM and SignalP6 (S3 Table). Abbreviations: IM: Inner membrane; LCV: Legionella-containig vacuole; OM: Outer membrane; TMS: Transmembrane segment.

## Materials and methods

### Materials

Chemicals were from Sigma-Aldrich unless otherwise specified. SERVAGel™ TG PRiME™ 8–16 % precast gels were from Serva. Primers were synthetized by Eurofins and Integrated DNA Technologies. Monoclonal anti-HA antibody was from Sigma-Aldrich. Secondary antibodies goat anti-mouse IgG DyLight 800 conjugate and goat anti-rabbit IgG DyLight 680 conjugate were from Thermo-Fisher (SA5-35571, 35568, respectively).

### Bacterial strains and plasmids

Bacterial strains and plasmids used in this study are listed in S4 Table. Primers for construction of strains and plasmids are also listed in S4 Table. All *Legionella* strains were derived from *Legionella pneumophila* strain Philadelphia-1 (LP01). Bacterial cultures were supplemented as required with chloramphenicol (6.8 μg/mL). Protein overexpression from the low-copy number plasmid pxDC61 was induced by supplementing the media with 0.5 mM Isopropyl-β-D-thiogalactopyranosid (IPTG). Molecular cloning was performed by standard Gibson cloning [38] using templates and primers as listed in Supplementary Data. pXDC61 was a gift from Howard Shuman (Addgene plasmid #21841; http://n2t.net/addgene:21841; RRID:Addgene_21841). The in-frame deletion of *dotG* into the *Legionella* strain LP01 was generated using allelic exchange as previously described [39]. Briefly, 1000 bp upstream and downstream to the *dotG* gene were cloned in the suicide vector pSR47S [40]. The deletion plasmid was transduced by mating from *Escherichia coli* CR14 into LP01 with the help of the *E. coli* strain CR19 [41].

### Immunoblotting

For protein detection, samples were subjected to SDS PAGE using SERVAGel™ TG PRiME™ 8–16% precast gels, transferred onto a PVDF membrane (Bio-Rad), and probed with primary antibodies anti-HA (1:1,000). Secondary antibodies were goat anti-rabbit IgG DyLight 800 conjugate (1:10,000). Scanning of the PVDF membrane and image analysis was performed with a Li-Cor Odyssey scanner and Image Studio 2.1.10 (Li-Cor).

### Crude membrane preparation

Crude membranes were prepared as reported previously [42]. 8 OD units of *L*. *pneumophila* cultures were resuspended in 750 μl buffer K (50 mM triethanolamine (TEA), pH 7.5, 250 mM sucrose, 1 mM EDTA, 1 mM MgCl_2_, 10 μg/ml DNAse I, 2 mg/ml lysozyme, 1:100 protease inhibitor cocktail). After incubation on ice for 30 min, samples were bead milled (SpeedMill Plus, Jena Analytics). Beads, unbroken cells and debris were removed by centrifugation for 20 min at 20,000 × *g* and 4 °C. Crude membranes were pelleted by centrifugation for 45 min at 186.007 x *g* and 4 °C in a Beckman TLA 55 rotor. Pellets containing crude membranes were frozen until use.

### Membrane fractionation

Inner and outer membranes were fractionated from 1000 OD units of *L. pneumophila* culture using a continuous sucrose gradient (30–50 %, w/w). In brief, 300 mL of a culture of *L. pneumophila* was lysed in buffer K by three cycles of French pressing at 20,000 psi. Debris were removed by centrifugation at 8,000 x g for 20 min at 4 °C. Crude membranes were precipitated at 235,000 × *g* for 45 min, resuspended in buffer M (50 mM TEA, pH 7.5, 1 mM EDTA), and loaded on top of a 30–50 % (w/w) continuous sucrose gradient, which was prepared using a Gradient Station (Biocomp, Fredericton, NB, Canada). Inner and outer membranes were separated by centrifugation at 285,000 × *g* for 15 h in a Beckman SW 41 swing out rotor. Twelve fractions of ∼1.1 ml were collected using the Gradient Station.

### Urea extraction

Membrane samples were solubilized in 8 M urea in 2× buffer M (100 mM TEA, pH 7.5, 2 mM EDTA) for 1 h at room temperature and centrifuged for 1.5 h at 23 °C in the Beckman TLA-55 rotor at 186.007 x *g* Pellets were resuspended in SB loading buffer, heated at 50 °C for 15 min, and analyzed by SDS PAGE, Western blotting, and immunodetection using anti HA antibody.

### Split Nanoluc-based translocation assay

HiBiT-tagged effector proteins were overexpressed in *L. pneumophila* using the plasmid pxDC61. One day prior to infection, RAW 264.7 macrophages expressing LgBiT [28,29] were seeded in a white clear-bottom 96 well cell culture plate (Greiner) at 8 x 10^4^ cells/well. *L. pneumophila* strains were grown for 2-3 days on BCYE-Agar plates supplemented with “BCYE Growth Supplements” (Oxoid) and appropriate antibiotics. Gene expression was induced by incubation of bacteria at 37 °C on BCYE agar plates with 0.5 mM IPTG for 24 hours. 100 µl of bacteria resuspended in HBSS + DrkBiT (1:1000) at 1.6×10^8^ cfu/ml were used to infect the RAW 264.7 macrophages (MOI = 200). After centrifugation at 300 x g for 8 minutes, 25 µl of NanoGlo® live cell buffer (Promega) supplemented with 1:20 of the extended live cell substrate Endurazine^TM^ (Promega) was added to each well. Luminescence was measured at 37°C and 5 % CO_2_ every 15 minutes for 12 hours in a Tecan Spark plate reader.

### Split Nanoluc-based topology assay

HiBiT-tagged effector proteins were overexpressed in *L. pneumophila* using the plasmid pxDC61. *L. pneumophila* strains were grown for 2-3 days on BCYE-Agar plates supplemented with “BCYE Growth Supplements” (Oxoid) and appropriate antibiotics. Gene expression was induced by incubation of bacteria at 37 °C on BCYE agar plates including 0.5 mM IPTG for 24 hours. 0.5 OD units of *L. pneumophila* were suspended in 500 µl PBS containing 10 μg/ml DNase I, 1 mM MgCl_2_, and 1:100 protease inhibitor cocktail. To obtain total luminescence 2 mg/ml lysozyme and 0.5 % (v/v) Triton-X 100 was added to the appropriate samples. All samples were incubated for 30 min at 4 °C. The luminescence was measured in a 384 well plate using NanoGlo® live cell buffer (Promega) supplemented with 1:50 of either a membrane impermeable substrate (Promega), for the periplasmic samples, or the lytic substrate (HiBiT lytic working solution, Promega), for the total luminescence samples.

### DeepTMHMM prediction

The DeepTMHMM web server (https://dtu.biolib.com/DeepTMHMM) in the version 1.0.24 (03/30/2023) was used to generate membrane topology predictions for all proteins of interest [24].

### Statistical analysis

All analyses were performed using R (R Core Team, 2017) with R commander [43]. The distribution of the data was assessed using a Shapiro-Wilk or a Kolmogorov-Smirnov statistic with Lilliefors correction. If the data were normally distributed, a Student’s t-test considering equality of variances was used to compare two groups, and a post-hoc Tukey’s honestly significant difference test was used to compare more than two groups pairwise. If the data were not normally distributed, a Wilcoxon test was used to compare two groups and a post-hoc Dunn test with Benjamini-Hochberg correction method was used to compare more than two groups pairwise. The significance level was set a priori at α = 0.05.

### NanoLC-MS/MS analysis

Double in-gel digests of proteins were performed with trypsin and chymotrypsin as described previously [44]. Extracted and desalted [45] peptides were analysed on an Easy-nLC 1200 system coupled to a Q Exactive HF mass spectrometer (all Thermo Fisher Scientific) [44]. In brief: peptides were separated by reversed phase chromatography using a 57 min or 87 min segmented gradient from 10-33-50-90% of HPLC solvent B (80% acetonitrile in 0.1% formic acid) in HPLC solvent A (0.1% formic acid) at a flow rate of 200 nl/min. The mass spectrometer was operated in data-dependent mode. Full scans were acquired in the mass range of 300-1650 *m/z* at a resolution of 60,000. Target values were set to 3×10^6^ charges with maximum IT set to 25 ms. The 7 or 12 most intense precursor ions were sequentially fragmented in each scan cycle using higher energy collisional dissociation (HCD). MS/MS spectra were recorded with a resolution of 60,000 or 30,000. AGC target was set to 10^5^, whereby fill times were set to 110 ms and 45 ms, respectively. In all measurements, sequenced precursor masses were excluded from further selection for 30 s.

### MS data processing

Processing of MS spectra was performed with MaxQuant software package version 1.5.2.8 [46] with integrated Andromeda search engine [47]. MS datasets were searched against a *Legionella pneumophila* subsp. Pneumophila (strain Philadelphia 1/ATCC33152/DSM7513) database obtained from Uniprot on 03/19/2018 (https://www.uniprot.org/<u>; 2,953 entries</u>), and against a database of 285 commonly observed contaminants. Settings were defined as described previously [44]: trypsin and chymotrypsin were defined as proteases, and a maximum of five missed cleavages were allowed. Oxidation of methionine, and protein N-terminal acetylation were chosen as variable modifications, and carbamidomethylation on cysteine was specified as fixed modification. Mass tolerance was set to 4.5 parts per million (ppm) for precursor ions and 20 ppm for fragment ions. Peptide, protein and modification site identifications were reported at a false discovery rate (FDR) of 0.01, estimated by the target-decoy approach [48]. The “match between runs” option was enabled, and iBAQ (Intensity Based Absolute Quantification) and LFQ (Label-Free Quantification) values were calculated [49]. Dependent on the dataset, the minimum ratio count was set to 1, and MS/MS was not required for LFQ comparison.

The MS data have been deposited to the ProteomeXchange Consortium (http://proteomecentral.proteomexchange.org) via the PRIDE partner repository with the data set identifier PXD050153.

## Acknowledgements

We thank Hiroki Nagai for providing published mutant strains of *L. pneumophila*. We thank Silke Wahl and Irina Droste-Borel for their support with sample preparation for MS measurements. We thank Andrea Eipper for technical support in the laboratory. We thank John Jairo Aquilera Correa for his support with the statistical analysis of the data.

## Supplemental Information

**S1 Table 3line file of the DeepTMHMM prediction of the 82 previously reported TMD-effectors**

**S2 Table gff3 file of the DeepTMHMM prediction of the 82 previously reported TMD-effectors**

**S3 Table DeepTMHMM prediction of the 82 previously reported TMD-effectors**

**S4 Table Primers, Plasmids and Strains S5 Table Statistics, p-values**

**S1 Fig.**
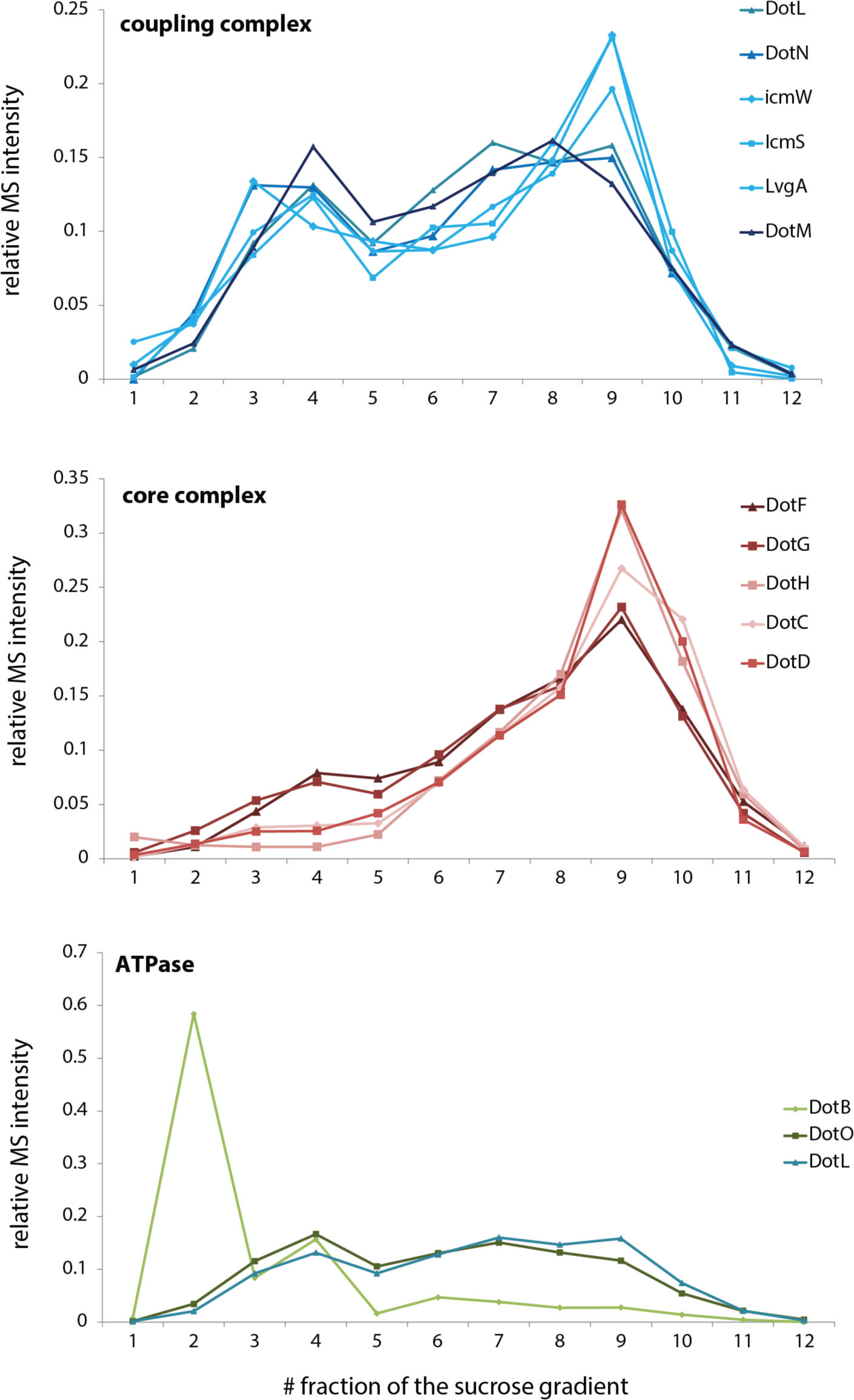
Distribution of T4bSS components across the membrane-fractionating sucrose gradient. The relative abundance of the indicated T4bSS components in 12 fractions of a membrane-fractionating sucrose gradient as analyzed by mass spectrometry. The coupling complex components as well as DotO and DotL follow the distribution of known inner membrane proteins (see Fig. 1A). DotB follows the distribution of soluble proteins and the core complex components follow the distribution of outer membrane proteins. The equilibration of the core complex in the outer membrane fractions may result from the high density of vesicles that contain the large core complex.

## References

1. Costa TRD, Felisberto-Rodrigues C, Meir A, Prevost MS, Redzej A, Trokter M, et al. Secretion systems in Gram-negative bacteria: structural and mechanistic insights. Nat Rev Micro. 2015;13: 343–359. doi:10.1038/nrmicro3456

2. Galán JE, Waksman G. Protein-injection machines in bacteria. Cell. 2018;172: 1306–1318. doi:10.1016/j.cell.2018.01.034

3. Nagai H, Roy CR. The DotA protein from Legionella pneumophila is secreted by a novel process that requires the Dot/Icm transporter. EMBO J. 2001;20: 5962–5970. doi:10.1093/emboj/20.21.5962

4. Qiu J, Luo Z-Q. Legionella and Coxiella effectors: strength in diversity and activity. Nat Rev Micro. 2017;15: 591–605. doi:10.1038/nrmicro.2017.67

5. Gomez-Valero L, Rusniok C, Carson D, Mondino S, Pérez-Cobas AE, Rolando M, et al. More than 18,000 effectors in the Legionella genus genome provide multiple, independent combinations for replication in human cells. Proc Natl Acad Sci. 2019;116: 2265–2273. doi:10.1073/pnas.1808016116

6. Durie CL, Sheedlo MJ, Chung JM, Byrne BG, Su M, Knight T, et al. Structural analysis of the Legionella pneumophila Dot/Icm type IV secretion system core complex. eLife. 2020;9: e59530. doi:10.7554/elife.59530

7. Meir A, Macé K, Lukoyanova N, Chetrit D, Hospenthal MK, Redzej A, et al. Mechanism of effector capture and delivery by the type IV secretion system from Legionella pneumophila. Nat Commun. 2020;11: 2864. doi:10.1038/s41467-020-16681-z

8. Sheedlo MJ, Ohi MD, Lacy DB, Cover TL. Molecular architecture of bacterial type IV secretion systems. Plos Pathog. 2022;18: e1010720. doi:10.1371/journal.ppat.1010720

9. Dutka P, Liu Y, Maggi S, Ghosal D, Wang J, Carter SD, et al. Structure and function of the Dot/Icm T4SS. bioRxiv. 2023; 2023.03.22.533729. doi:10.1101/2023.03.22.533729

10. Kitao T, Kubori T, Nagai H. Recent advances in structural studies of the Legionella pneumophila Dot/Icm type IV secretion system. Microbiol Immunol. 2022;66: 67–74. doi:10.1111/1348-0421.12951

11. Luo Z-Q, Isberg RR. Multiple substrates of the Legionella pneumophila Dot/Icm system identified by interbacterial protein transfer. Proc Natl Acad Sci. 2004;101: 841–846. doi:10.1073/pnas.0304916101

12. Nagai H, Cambronne ED, Kagan JC, Amor JC, Kahn RA, Roy CR. A C-terminal translocation signal required for Dot/Icm-dependent delivery of the Legionella RalF protein to host cells. Proc Natl Acad Sci USA. 2005;102: 826–831. doi:10.1073/pnas.0406239101

13. Jeong KC, Sutherland MC, Vogel JP. Novel export control of a Legionella Dot/Icm substrate is mediated by dual, independent signal sequences. Mol Microbiol. 2015;96: 175–188. doi:10.1111/mmi.12928

14. Huang L, Boyd D, Amyot WM, Hempstead AD, Luo Z-Q, O’Connor TJ, et al. The E Block motif is associated with Legionella pneumophila translocated substrates. Cell Microbiol. 2011;13: 227–245. doi:10.1111/j.1462-5822.2010.01531.x

15. Kwak M-J, Kim JD, Kim H, Kim C, Bowman JW, Kim S, et al. Architecture of the type IV coupling protein complex of Legionella pneumophila. Nat Microbiol. 2017;2: 17114. doi:10.1038/nmicrobiol.2017.114

16. Meir A, Chetrit D, Liu L, Roy CR, Waksman G. Legionella DotM structure reveals a role in effector recruiting to the Type 4B secretion system. Nat Commun. 2018;9: 507. doi:10.1038/s41467-017-02578-x

17. Dolezal P, Aili M, Tong J, Jiang J-H, Marobbio CMT, Marobbio CM, et al. Legionella pneumophila secretes a mitochondrial carrier protein during infection. Roy CR, editor. PLoS Pathog. 2012;8: e1002459. doi:10.1371/journal.ppat.1002459

18. Weigele BA, Orchard RC, Jimenez A, Cox GW, Alto NM. A systematic exploration of the interactions between bacterial effector proteins and host cell membranes. Nat Commun. 2017;8: 532. doi:10.1038/s41467-017-00700-7

19. Krampen L, Malmsheimer S, Grin I, Trunk T, Lührmann A, Gier J-W de, et al. Revealing the mechanisms of membrane protein export by virulence-associated bacterial secretion systems. Nat Commun. 2018;9: 3467. doi:10.1038/s41467-018-05969-w

20. Schibich D, Gloge F, Pöhner I, Björkholm P, Wade RC, Heijne GV, et al. Global profiling of SRP interaction with nascent polypeptides. Nature. 2016;536: 219–223. doi:10.1038/nature19070

21. Wagner S, Baars L, Ytterberg AJ, Klussmeier A, Wagner CS, Nord O, et al. Consequences of membrane protein overexpression in Escherichia coli. Mol Cell Proteomics. 2007;6: 1527–1550. doi:10.1074/mcp.m600431-mcp200

22. Wagner S, Königsmaier L, Lara-Tejero M, Lefebre M, Marlovits TC, Galán JE. Organization and coordinated assembly of the type III secretion export apparatus. Proc Natl Acad Sci USA. 2010;107: 17745–17750. doi:10.1073/pnas.1008053107

23. Bi D, Liu L, Tai C, Deng Z, Rajakumar K, Ou H-Y. SecReT4: a web-based bacterial type IV secretion system resource. Nucleic Acids Res. 2013;41: D660–D665. doi:10.1093/nar/gks1248

24. Hallgren J, Tsirigos KD, Pedersen MD, Armenteros JJA, Marcatili P, Nielsen H, et al. DeepTMHMM predicts alpha and beta transmembrane proteins using deep neural networks. 2022. doi:10.1101/2022.04.08.487609

25. Dixon AS, Schwinn MK, Hall MP, Zimmerman K, Otto P, Lubben TH, et al. NanoLuc complementation reporter optimized for accurate measurement of protein interactions in cells. ACS Chem Biol. 2016;11: 400–408. doi:10.1021/acschembio.5b00753

26. Heijne G von. Control of topology and mode of assembly of a polytopic membrane protein by positively charged residues. Nature. 1989;341: 456–458. doi:10.1038/341456a0

27. Kumazaki K, Chiba S, Takemoto M, Furukawa A, Nishiyama K-I, Sugano Y, et al. Structural basis of Sec-independent membrane protein insertion by YidC. Nature. 2014;509: 516–520. doi:10.1038/nature13167

28. Westerhausen S, Nowak M, Torres-Vargas CE, Bilitewski U, Bohn E, Grin I, et al. A NanoLuc luciferase-based assay enabling the real-time analysis of protein secretion and injection by bacterial type III secretion systems. Mol Microbiol. 2020;113: 1240–1254. doi:10.1111/mmi.14490

29. Fromm K, Boegli A, Ortelli M, Wagner A, Bohn E, Malmsheimer S, et al. Bartonella taylorii: A model organism for studying Bartonella infection in vitro and in vivo. Front Microbiol. 2022;13: 913434. doi:10.3389/fmicb.2022.913434

30. Herbel SM, Moyon L, Christ M, Elsayed EM, Caffrey BE, Malmsheimer S, et al. Screening for eukaryotic motifs in Legionella pneumophila reveals Smh1 as bacterial deacetylase of host histones. Virulence. 2022;13: 2042–2058. doi:10.1080/21505594.2022.2149973

31. Yamamoto M, Du Q, Song J, Wang H, Watanabe A, Tanaka Y, et al. Cell–cell and virus–cell fusion assay–based analyses of alanine insertion mutants in the distal α9 portion of the JRFL gp41 subunit from HIV-1. J Biol Chem. 2019;294: 5677–5687. doi:10.1074/jbc.ra118.004579

32. Amyot WM, deJesus D, Isberg RR. Poison domains block transit of translocated substrates via the Legionella pneumophila Icm/Dot System. Infect Immun. 2013;81: 3239–3252. doi:10.1128/iai.00552-13

33. Radics J, Königsmaier L, Marlovits TC. Structure of a pathogenic type 3 secretion system in action. Nat Struct Mol Biol. 2014;21: 82–87. doi:10.1038/nsmb.2722

34. Jong MFD, Sun Y, Hartigh ABD, Dijl JMV, Tsolis RM. Identification of VceA and VceC, two members of the VjbR regulon that are translocated into macrophages by the Brucella type IV secretion system. Mol Microbiol. 2008;70:<otherinfo> 13</otherinfo>78–1396. doi:10.1111/j.1365-2958.2008.06487.x

35. Locht C, Coutte L, Mielcarek N. The ins and outs of pertussis toxin. FEBS J. 2011;278: 4668– 4682. doi:10.1111/j.1742-4658.2011.08237.x

36. Barsy M de, Jamet A, Filopon D, Nicolas C, Laloux G, Rual J, et al. Identification of a Brucella spp. secreted effector specifically interacting with human small GTPase Rab2. Cell Microbiol. 2011;13: 1044–1058. doi:10.1111/j.1462-5822.2011.01601.x

37. Ninio S, Zuckman-Cholon DM, Cambronne ED, Roy CR. The Legionella IcmS-IcmW protein complex is important for Dot/Icm-mediated protein translocation. Mol Microbiol. 2005;55: 912–26. doi:10.1111/j.1365-2958.2004.04435.x

38. Gibson DG, Young L, Chuang R-Y, Venter JC, Hutchison CA, Smith HO. Enzymatic assembly of DNA molecules up to several hundred kilobases. Nat Methods. 2009;6: 343–345. doi:10.1038/nmeth.1318

39. Zuckman DM, Hung JB, Roy CR. Pore-forming activity is not sufficient for Legionella pneumophila phagosome trafficking and intracellular growth. Mol Microbiol. 1999;32: 990–1001. doi:10.1046/j.1365-2958.1999.01410.x

40. Merriam JJ, Mathur R, Maxfield-Boumil R, Isberg RR. Analysis of the Legionella pneumophila fliI gene: intracellular growth of a defined mutant defective for flagellum biosynthesis. Infection and Immunity. 1997;65: 2497–2501.

41. Roy CR, Isberg RR. Topology of Legionella pneumophila DotA: an inner membrane protein required for replication in macrophages. Infection and Immunity. 1997;65: 571–578. Available: /pmc/articles/PMC176098/?report=abstract

42. Dietsche T, Mebrhatu MT, Brunner MJ, Abrusci P, Yan J, Franz-Wachtel M, et al. Structural and functional characterization of the bacterial type III secretion export apparatus. Coombes BK, editor. PLoS Pathog. 2016;12: e1006071. doi:10.1371/journal.ppat.1006071

43. Fox J. The R Commander: A Basic-Statistics Graphical User Interface to R. J Stat Softw. 2005;14. doi:10.18637/jss.v014.i09

44. Singh N, Kronenberger T, Eipper A, Weichel F, Franz-Wachtel M, Macek B, et al. Conserved salt bridges facilitate assembly of the helical core export apparatus of a Salmonella enterica type III secretion system. J Mol Biol. 2021;433: 167175. doi:10.1016/j.jmb.2021.167175

45. Rappsilber J, Mann M, Ishihama Y. Protocol for micro-purification, enrichment, pre-fractionation and storage of peptides for proteomics using StageTips. Nat Protoc. 2007;2: 1896–1906. doi:10.1038/nprot.2007.261

46. Cox J, Mann M. MaxQuant enables high peptide identification rates, individualized p.p.b.-range mass accuracies and proteome-wide protein quantification. Nat Biotechnol. 2008;26: 1367–1372. doi:10.1038/nbt.1511

47. Cox J, Neuhauser N, Michalski A, Scheltema RA, Olsen JV, Mann M. Andromeda: a peptide search engine integrated into the MaxQuant environment. J Proteome Res. 2011;10: 1794–1805. doi:10.1021/pr101065j

48. Elias JE, Gygi SP. Target-decoy search strategy for increased confidence in large-scale protein identifications by mass spectrometry. Nat Methods. 2007;4: 207–214. doi:10.1038/nmeth1019

49. Tyanova S, Temu T, Cox J. The MaxQuant computational platform for mass spectrometry-based shotgun proteomics. Nat Protoc. 2016;11: 2301–2319. doi:10.1038/nprot.2016.136

